# A transfer-learning approach for first-year developmental infant brain segmentation using deep neural networks

**DOI:** 10.1101/2020.05.22.110619

**Authors:** Yun Wang, Fateme Sadat Haghpanah, Natalie Aw, Andrew Laine, Jonathan Posner

## Abstract

The months between birth and age 2 are increasingly recognized as a period critical for neuro-development, with potentially life-long implications for cognitive functioning. However, little is known about the growth trajectories of brain structure and function across this time period. This is in large part because of insufficient approaches to analyze infant MRI scans at different months, especially brain segmentation. Addressing technical gaps in infant brain segmentation would significantly improve our capacity to efficiently measure and identify relevant infant brain structures and connectivity, and their role in long-term development. In this paper, we propose a transfer-learning approach based on convolutional neural network (CNN)-based image segmentation architecture, QuickNAT, to segment brain structures for newborns and 6-month infants separately. We pre-trained QuickNAT on auxiliary labels from a large-scale dataset, fine-tuned on manual labels, and then cross-validated the model’s performance on two separate datasets. Compared to other commonly used methods, our transfer-learning approach showed superior segmentation performance on both newborns and 6-month infants. Moreover, we demonstrated improved hippocampus segmentation performance via our approach in preterm infants.

## 1 Introduction

Human brain development from birth to ∼2 years of age is dynamic and increasingly recognized as crucial for establishing cognitive abilities and behaviors that last a lifetime, as well as risk for neuro-psychiatric disorders, e.g., autism and schizophrenia [1]. Thus, it is crucial to understand and quantify typical neurodevelopmental trajectories from which deviant maturation indicative of developmental delay and/or disorders can be identified, as early and precisely as possible. Despite their importance, little is known about trajectories of structural and functional brain development during this sensitive period, and even less is known about how deviations in these trajectories relate to emerging cognition and behavior or predict later developmental outcomes. This significant scientific gap is partially due to current technical limits on the rigorous and valid quantification of brain structure and functioning in infants via MRI. MRI is an important, non-invasive approach for the study of developmental neuroscience. However, there are insufficient approaches available for accurate and automated (i.e., time-efficient) segmentation of early brain structural MRIs - a process that is vital for virtually all quantitative analyses across MRI modalities, e.g., diffusion MRI, functional MRI. Without accurate and automated segmentation, infant MRI analysis is prone to systematic errors and is labor-intensive, limiting both sample sizes, reliability, and reproducibility. Addressing the challenges of first-year developmental infant brain segmentation will significantly advance efforts toward early identification of developmental delays and/or developmental disorders and monitoring the effects of interventions.

As opposed to MRI scans from adults for which various automated pipelines are effective (e.g., FAST[2], SPM, FreeSurfer[3]), in infants, the segmentation step is by far the most difficult to implement. Compared with adults, infant brains 1) have a much lower contrast-to-noise ratio due to the relative lack of myelination and shorter scan times; 2) lower resolution due to the smaller overall volume of the brain; 3) reversal of gray matter (GM)/ white matter (WM) voxel intensity values before 6 months of age due to the transiently lower intensity of unmyelinated WM fibers relative to GM; and

4) similar GM/WM intensity values between 6-9 months, which reach adult values by 1 year of age. Another difficulty in applying a standardized approach is that the early infant brain experiences dramatic shifts in brain development over months. This creates difficulty in adapting extant infant tissue atlases, generally limited to a single age, to guide segmentation of scans acquired at other, even proximal ages.

In response to these challenges, various methods and pipelines have been attempted to segment the early infant brain MRI scans, e.g., DrawEM[4], iBEAT[5], MANTiS[6]. However, the smooth integration of these methods into a typical infant research workflow has been found challenging in different aspects. First, these methods often only work at fixed ages, such as newborns and 1 year old. Second, these methods are computationally demanding and time-consuming. Lastly, good comparability with manual segmentation by an expert neuroanatomist has yet to be achieved when applying these methods into an independent infant MRI dataset.

Convolutional neural network (CNN)-based methods have been introduced for infant brain segmentation in the last few years. In the most recent six-month infant brain segmentation challenge, the top-ranked team achieved good segmentation accuracy via adapted CNN architectures[7]. However, the number of training and testing subjects are small (N=10); therefore, their trained models may not be applicable to the infant scans acquired from different ages, sites, scanners or imaging protocols based on limited annotated data. A recent adaptation of a deep CNN called QuickNAT [8] has been documented to yield good accuracy and reproducibility on a large variety of adult neuroimaging datasets with 20-seconds-per-subject speed.

In this paper, we propose to pre-train QuickNAT on a large-scale neonatal MRI dataset with auxiliary labels (***Dataset I***: n=473; ages 0.8 ± 0.7 months) generated from existing neonatal brain segmentation software. Subsequently, in a transfer-learning approach, we fine-tune and cross-validate the pretrained model on manual labels from two independent infant MRI datasets with significant different age distributions (***Dataset II***: n=10; ages 1.1 ± 0.2 months; ***Dataset III***: n=10; ages 6 ± 0.5 months). Third, we assess the segmentation performance of our strategy compared with other commonly used approaches in term-born infant and preterm infants.

## 2 Materials and Methods

### 2.1 Imaging Datasets

We used three independent, publicly available infant datasets to train and test our deep learning models. Of note, these datasets come from different MRI scanners and used different imaging protocols (Table 1), which we leveraged to optimize the training and testing of the model. We resampled MRI scans from ***Dataset II*** and ***Dataset III*** to match the imaging dimension and voxel resolutions of Dataset I. Because of the higher tissue contrast in T2-weighted images, we only used T2w structural scans for the automatic segmentation.

**Table 1.** Demographics, MRI Imaging Protocols, and Labels for Three Datasets

|  |  | *Dataset I* | *Dataset II* | *Dataset III* |
| --- | --- | --- | --- | --- |
| Subject Number |  | 473 | 10 | 10 |
| Age at scan (months) | | $0.8 \pm 0.7$ | $1.1 \pm 0.2$ | $6.0 \pm 0.5$ |
| Gender |  |  |  |  |
|  | F | 207 | 4 | 5 |
|  | M | 266 | 6 | 5 |
| MRI Scanners |  | 3T Philips | 3T Siemens | 3T Siemens |
| MRI Image Dimensions |  | 290×290×203 | 304×304×157 | 144×192×256 |
| MRI Voxel Resolution |  | 0.5×0.5×0.5mm | 0.63×0.63×0.63mm | 1×1×1mm |
| Training/Test Slices |  | **96,019** | 1,570 | 2,560 |

Ages are presented as mean ± standard deviation.

***Dataset I*: Developing Human Connectome Project (dHCP)** comes from an ongoing large-scale collaborative project. We used 473 subjects from the second dHCP data release. More details about the imaging parameters and preprocessing procedures can be found in the original work [9]. Auxiliary labels (87 regions and 9 tissues) used for the pre-training CNN model were generated by the dHCP structural minimal processing pipeline.

***Dataset II*: Melbourne Children’s Regional Infant Brain (M-CRIB)**, consists of 10 neonatal subjects. Imaging parameters and preprocessing procedures are described in [6]. The M-CRIB dataset was manually annotated with 100 regions labels by an experienced neuroanatomist.

***Dataset III*: iSeg-2017** is a subset of the 2017 6-month infant MRI brain segmentation MICCAI challenge dataset. Imaging parameters and preprocessing procedures can be found in [7]. Manually annotated 3 tissue labels were released along with this dataset.

### 2.2 QuickNAT Architecture

QuickNAT is based on a fully convolutional neural network for the segmentation of whole-brain neuroanatomy via adult T1-weighted MRI scans. Technically, QuickNAT is a modified version of U-Net with skip connections, enhanced by unpooling layers, which makes the architecture more capable of segmenting small subcortical structures. Also, QuickNAT introduces dense connections in both encoder and decoder blocks to aid gradient flow from the shallower to deeper layers and to promote feature re-usability. This is essential given the limited amount of training data in the field of infant brain research. To simplify our work, in this paper, we only used the axial slices of two-dimensional (2D) T2 weighted MRI data as input images and output segmented images with multi-class labels (including tissues and anatomical regions). We chose multi-class segmentation classifiers over binary classifiers because we aimed to evaluate the model’s performance for differentiating adjacent tissues. A detailed explanation of QuickNAT can be found in [8].

### 2.3 Performance Analysis

In this paper, we ran three deep learning experiments with hyperparameter fine-tuning and segmentation performance evaluation. We also made a comparison between our transfer-learning approach and other commonly used approaches.

- **Cross-Validation**: In experiment 1, we validated our model in the holdout 20% ***Dataset I***. In experiments 2 and 3, we performed a leave-one-out cross-validation across the entire *Datasets II* and *III*.
- **Hyperparameter fine-tuning**: We trained each model for 50 epochs with 9 different pre-defined learning rates (see supplemental materials) and chose the optimal learning rate based on segmentation accuracy on the test dataset. Batch size is set to 8, limited by the 24GB RAM of NVIDIA Geforce Titan RTX GPU.
- **Dice Score and Accuracy**: Model performance was evaluated with both accuracy and Dice Score. For accuracy, we calculated the intersection area of true labels (A) and predicted labels (B) divided by the area of the predicted label [Equation 1], where A and B denote the segmentation labels generated manually and computationally, respectively. Dice Score is 2 * the area of intersection of the true labels and predicted labels divided by the sum of the area of predicted labels and true labels [Equation 2]. Since the models are multi-class, we did the same calculation for each label in each experiment.

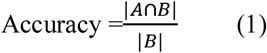

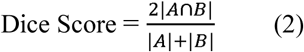

#### Other common segmentation approaches

In addition, we compared segmentation accuracy derived from our transfer-learning segmentation approach with other commonly used infant segmentation methods, including DrawEM, iBEAT, MANTiS, and FAST.

## 3 Experiments and Results

### 3.1 Experiment 1: Baseline model on *Dataset I* with auxiliary labels

In this experiment, we randomly split ***Dataset I*** into two parts: 80% of subjects with 2D axial slices for training (76,815) and the other 20% subjects (19,204) for testing the performance of the model. We trained this baseline model with hyperparameter tuning, including learning rates, epochs, and batch sizes. The optimal learning rate is 5×10^−5^. We tested the model’s performance by calculating the Dice Score and the accuracy on the test dataset for 87 regions and 9 different types of tissues separately (see supplemental materials). The goal of experiment 1 was to output an optimized baseline model with pre-trained weights. Our baseline model showed an average 88% accuracy and Dice Score for region segmentation and 95% accuracy and Dice Score on tissue segmentation (Table 2).

**Table 2.** Baseline Model Performance from Experiment 1

|  | Multi-region<br>Segmentation<br>(N=87) | Multi-tissue<br>Segmentation<br>(N=9) |
| --- | --- | --- |
| Mean Dice Score | $0.88 \pm 0.05$ | $0.95 \pm 0.03$ |
| Mean Accuracy | $0.88 \pm 0.06$ | $0.95 \pm 0.03$ |

Values are presented as mean ± standard deviation.

### 3.2 Experiment 2: Transfer pre-trained baseline model to *Dataset II*

Motivated by the superior performance of our baseline model (see 3.1), we applied the baseline model to ***Dataset II***, which has a similar age distribution as ***Dataset I***. We trained the model in two stages. In stage I, we took the initial weights of the baseline model from experiment 1 and modified the classifier in last layer to make the architecture compatible with the number of tissue/region labels in ***Dataset II.*** Then, we unfroze the last classifier layer and ran it for 15 epochs. In stage II, we unfroze the remaining layers and trained the entire model for 50 epochs. The optimal learning rate is 1×10^−4^, and details can be found in supplemental materials.

The confusion matrix is provided in Fig.1A. Overall, predicted labels from the proposed approach achieves 90% Dice Score. For each tissue, the mean accuracy of each epoch is calculated across all subjects after 10 iterations. Mean accuracy over epochs is plotted in Fig.1B, and no overfitting sign is observed. Of note, our transfer learning approach is capable of segmenting small subcortical structures such as the hippocampus in term-born infants, shown in Fig.1C.

**Fig. 1.**
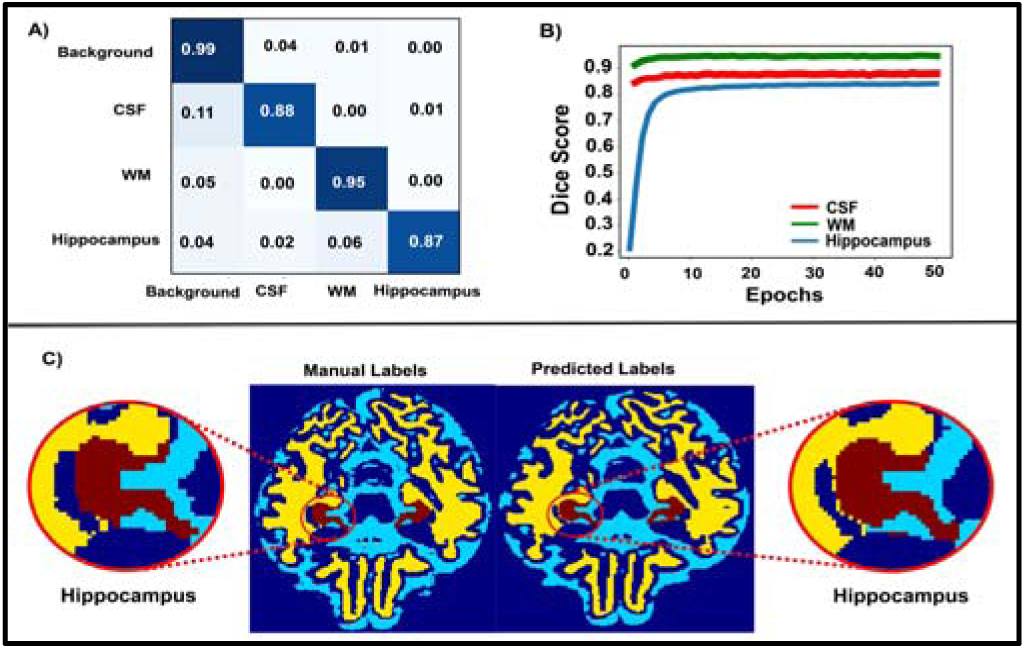
Transfer learning approach on multi-class tissue segmentation for ***Dataset II***. A) confusion matrix, B) mean accuracy over epochs for each tissue, C) A zoom view of hippocampus segmentation from our approach over manual labels.

### 3.3 Experiment 3: Transfer pre-trained baseline model to *Dataset III*

To test our proposed approach on segmenting infant brain structures across difference ages, we adapted similar training and testing procedures as experiment 2 on ***Dataset III***, which has an average age of six months. The optimal learning rate is 1×10^−4^, and details can be found in supplemental materials. The confusion matrix is provided in Fig.2A. Overall, predicted labels from our proposed approach achieves 85% Dice Score. Confusion occurs most for adjacent classes of WM and GM. For each tissue, the mean accuracy of each epoch is calculated across all subjects after 10 iterations. Mean accuracy over epochs is plotted in Fig.2B, and no overfitting sign is observed. We also inspected the performance of segmenting sulcal area (red arrow) and found good results (Fig.2C).

**Fig. 2.**
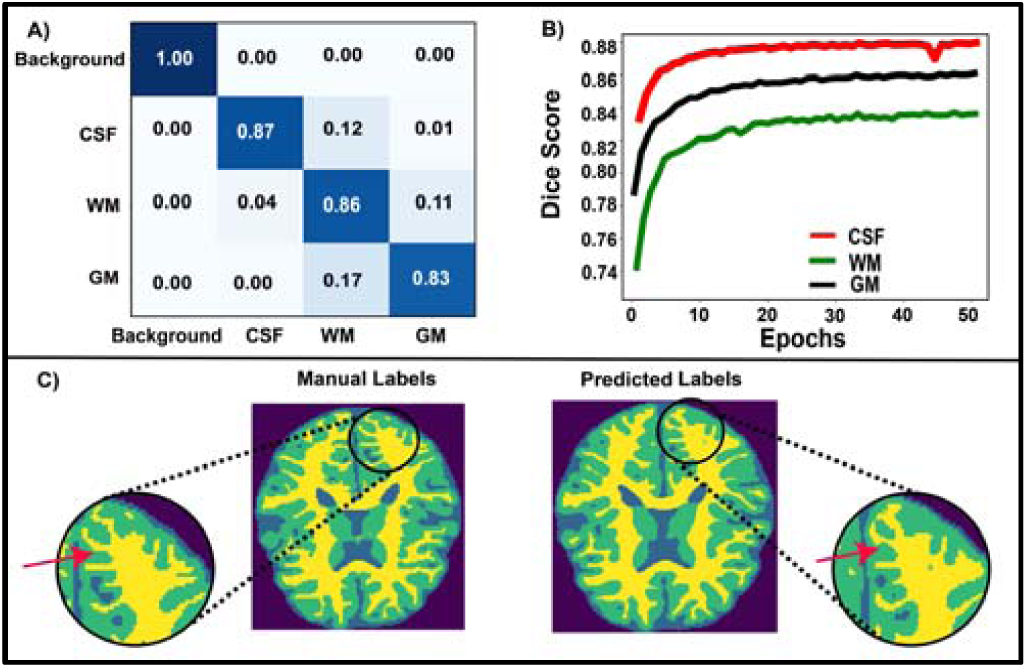
Transfer learning approach to six-month infant brain segmentation (***Dataset III***). A) confusion matrix, B) mean accuracy over epochs for each tissue, C) A zoom view shows the comparable performance of our proposed in segmenting sulci with manual labeling.

### 3.4 Comparing with other commonly used methods

For both ***Dataset II*** and ***Dataset III***, we calculated and compared segmentation performance as well as the computational speed by our transfer learning approaches with other methods, e.g., DrawEM, iBeat, MANTiS, and FAST.

Relative to other commonly used approaches or pipelines, our transfer learning approaches achieves highest Dice Score on both newborn (Fig.3A) and 6-month infant (Fig.3B) brain segmentation. In addition, our approach segments whole brain in 4 seconds, which is an order of magnitude faster than other approaches, see Fig. 3C.

**Fig. 3.**
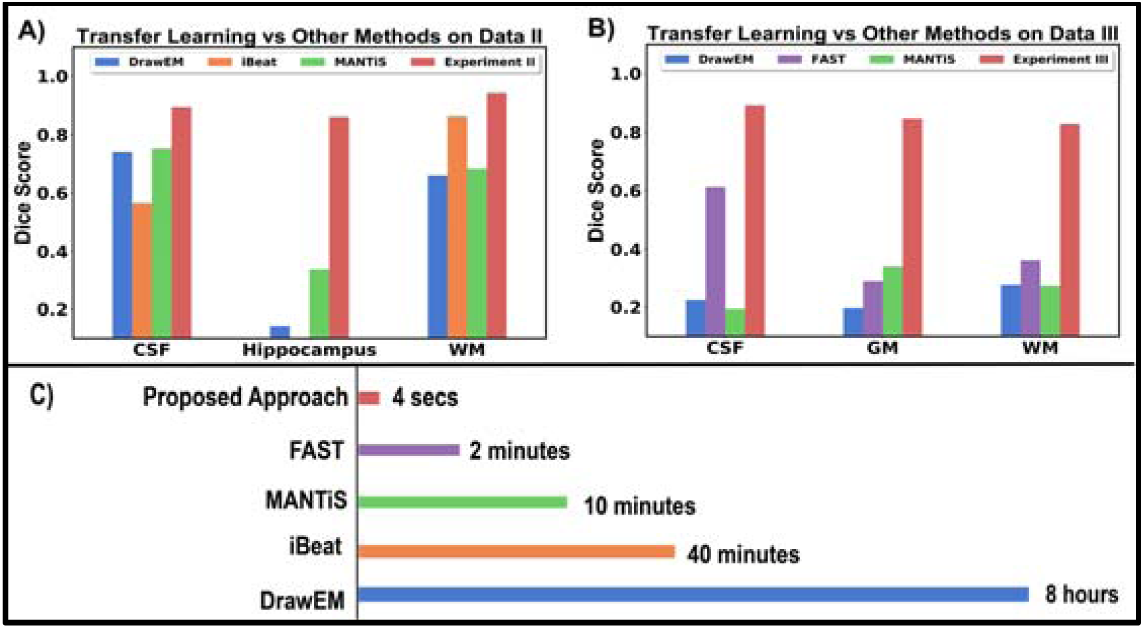
Segmentation performance comparisons on A) ***Dataset II***, B) ***Dataset III***. C) Segmentation speed for our proposed approach, FAST, MANTiS, iBeat, and DrawEM.

### 3.5 Clinical and research utility for precise hippocampus segmentation in preterm infants

In the above experiments, we observed that our approach provide accurate segmentation of the infant hippocampus. This could have significant value in clinical research because the hippocampus is a structure critical for learning, memory, and emotion regulation[10]. Moreover, atypical development of the hippocampus is posited to be related to several neuropsychiatric disorders including ADHD[11], schizophrenia[12], depression[13], and anxiety[14]. Lastly, the hippocampus is sensitive to early environmental insults such as preterm delivery[15, 16], where early moritoring of hippocampal maturation could help inform future interventions and precision medicine[17, 18]. To further explore this possibility, we investigated hippocampal segmentation accuracy via our transfer-learning approach in an infant prematurely born at 34 weeks obtained from the dHCP dataset. As shown in Fig. 4, our method provided greater accuracy than DrawEM segmentation.

**Fig. 4.**
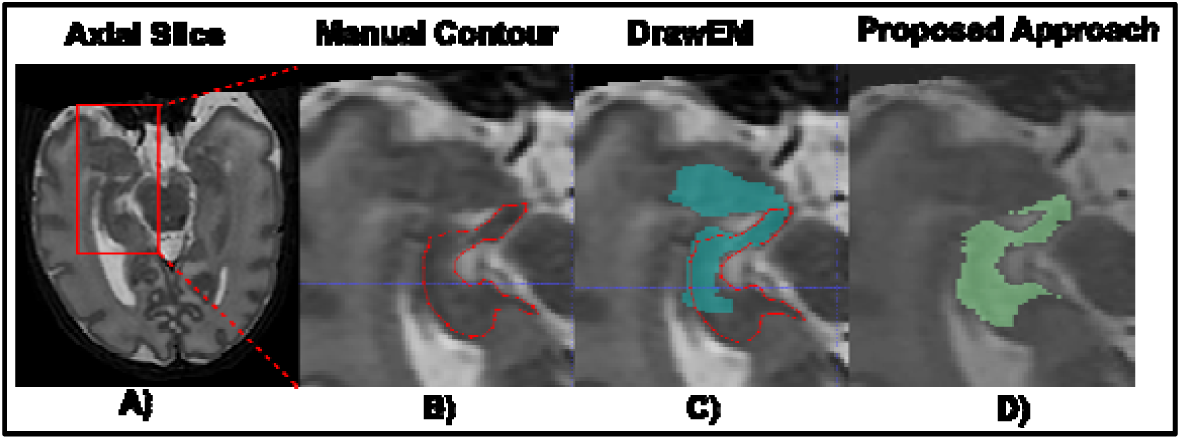
Hippocampus segmentation in one preterm infant. A) the 82nd axial slice, B) manual polygon red contour with ITK-SNAP around left hippocampus, C) DrawEM segmentation (blue color), D) segmentation from our transfer learning approach (green color).

## 4 Discussion

In this paper, we proposed a transfer-learning approach based on the QuickNAT architecture, which can transfer the knowledge of auxiliary labels learned from a large-scale public dataset to better segment brain neuroanatomy for infants at different ages from independent sources. Compared to other commonly used infant MRI processing methods, our approach significantly improved the accuracy and speed of segmenting brain structures for newborns and 6-month old infants, suggesting the potential for extending our work to diverse ages of infants. Moreover, via our transfer-learning approach, hippocampus segmentation accuracies improved in both preterm and term-born infants, indicating potential clinical and research utility for studying hippocampus development in prematurely born infants.

## Acknowledgments

This work was supported by Environmental Influences on Child Health Outcomes (ECHO) program (UG3OD023328, Jonathan Posner) as well as ECHO-Opportunities and Infrastructure Fund (Yun Wang). In addition, we want to remember 176 passengers who lost their precious lives in the tragic and unforgettable airplane crash of the Ukrainian International Flight, PS752. For our beloved friends, Pouneh Gorji and Arash Pourzarabi, two of the kindest souls living on the earth.

